# Relating excitatory and inhibitory neurochemicals to visual perception: a magnetic resonance study of occipital cortex between migraine events

**DOI:** 10.1101/477935

**Authors:** Yu Man Chan, Kabilan Pitchaimuthu, Qi-Zhu Wu, Olivia L Carter, Gary F Egan, David R Badcock, *Allison M McKendrick

**Affiliations:** Department of Optometry & Vision Sciences, The University of Melbourne, Melbourne, Victoria, Australia; Monash Biomedical Imaging, Monash University, Melbourne, Victoria, Australia; Shenzhen Sinorad Medical Electronics Inc., Shenzhen, China; Melbourne School of Psychological Sciences, University of Melbourne, Melbourne, Victoria, Australia; School of Psychological Science, University of Western Australia, Perth, Western Australia, Australia

## Abstract

Certain perceptual measures have been proposed as indirect assays of brain neurochemical status in people with migraine. One such measure is binocular rivalry, however, previous studies have not measured rivalry characteristics and brain neurochemistry together in people with migraine. This study compared spectroscopy-measured levels of GABA and Glx (glutamine and glutamate complex) in visual cortex between 16 people with migraine and 16 non-headache controls, and assessed whether the concentration of these neurochemicals explains, at least partially, inter-individual variability in binocular rivalry perceptual measures. Mean Glx level was significantly reduced in migraineurs relative to controls, whereas mean occipital GABA levels were similar between groups. Neither GABA levels, nor Glx levels correlated with rivalry percept duration. Our results thus suggest that the previously suggested relationship between rivalry percept duration and GABAergic inhibitory neurotransmitter concentration in visual cortex is not strong enough to enable rivalry percept duration to be reliably assumed to be a surrogate for GABA concentration, at least in the context of healthy individuals and those that experience migraine.

## 1 Introduction

Migraine is a very common neurological disorder that affects approximately 10-15% of the adult population (1). Migraine often involves the visual pathways. In approximately 30% of people with migraine, this involvement manifests as a visual aura (2). Other very common visual symptoms are photophobia, blur and visual discomfort. The common involvement of visual system in migraine symptomatology has resulted in extensive research that has used the visual system as a model for exploring the migrainous brain more generally (for review see: (3-5)). While there is still significant debate about the exact pathophysiology of migraine, convergent evidence from brain imaging and electrophysiology points to imbalanced cortical inhibition-excitation around the time of transient migraine events (6). 1H-magnetic resonance spectroscopy (1H-MRS) has been used to demonstrate altered cortical metabolite levels when people were relatively asymptomatic (interictally) (7-11), including reduced levels of inhibitory neurotransmitter GABA (gamma-Aminobutyric acid) in the occipital cortex (10), especially in those with recent and more severe migraines (11). Most studies report elevated glutamate (main excitatory neurotransmitter) interictally (12, 13) although not universally (10, 14). Abnormalities in the serotonergic system have also been implicated in migraine, once again with some equivocal results (15-19).

Perceptual studies also provide evidence for interictal imbalance between cortical inhibition and excitation in migraine (20, 21). Based on convergent evidence from primate neurophysiology, human brain imaging, and human behavioural studies, there are a number of perceptual tasks where performance is predicted to be altered by an imbalance between excitation and inhibition. Specifically, in the context of migraine, differences in perceptual performance have been found for centre-surround tasks (22, 23), for the motion after-effect (21, 24), and for the tilt illusion and tilt aftereffect (24, 25). Of particular relevance to this study is a previous observation that perceptual rivalry modulates more slowly in both auditory and visual domains in migraineurs (26, 27). Binocular rivalry is a specific form of perceptual rivalry that arises when different images are presented to the two eyes and compete for perceptual dominance (28-30). A series of recent neuroimaging studies have demonstrated that GABA concentration in visual cortex is correlated with perceived binocular rivalry switch rate (31-33), however these studies have not been conducted in migraine groups. The observed correlation between rivalry percept duration and GABA concentration is modest, with approximately 25% of the variance in switch rate explained by GABA concentration (33).

If perceptual measures can indeed provide reasonable correlates of brain neurochemical status, then it is possible that visual or auditory tests might be useful for monitoring of brain status in migraine, or perhaps even to predict migraine events (23). Perceptual testing is easy to perform, inexpensive, quick, and could potentially be undertaken on a daily basis throughout the migraine cycle in order to understand individual patterns of brain status. However, for perceptual rivalry to be useful as an indirect measure of the inhibitory-excitatory balance in migraine, a better understanding of the underlying neurochemical influences is required. It is overly simplistic to suggest that GABA is the only neurochemical governing perceptual binocular rivalry. Indeed binocular rivalry percept duration is also influenced by modulating the serotonergic pathways (34, 35). Given that both the GABA-ergic and Glutaminergic pathways, in addition to the serotonergic pathways have been previously implicated in migraine pathophysiology, it is not readily predictable whether perceptual rivalry switch rate should relate to GABA concentration in people with migraine.

Hence, in this study, we measured key excitatory and inhibitory neurochemicals in visual cortex in people with migraine between attacks using 1H-MRS (36), and related these to binocular rivalry perceptual data collected within approximately 1 hour of the neuroimaging. The aim of this study was to investigate if the concentration of GABA and the main cortical excitatory neurotransmitter complex, Glx (glutamine and glutamate complex) differs between groups, whether the ratio of these differs relative to non-headache controls, and whether concentration of these neurochemicals does indeed explain any of the inter-individual variability in binocular rivalry percept durations.

## 2 Methods

### 2.1 Participants

Sixteen non-headache controls (20 – 34 years; mean age 27.1 years; 8 males), nine migraine sufferers with aura (21 – 42 years; mean age 31.0 years; 1 male) and seven migraine sufferers without aura (20–49 years; mean age 31.1 years; 2 males) participated in this study. The age distribution of the controls were not statistically different from the combined migraine cohort (Independent samples t-test: t(15)=1.66,p=0.12). Sample size was determined based on previous work assessing similar types of cortical (7, 11, 12) and perceptual measures (26, 27) that have found significant outcomes based on 9 to 20 observers in each group. The total sample size of 32 allows for a correlation of 0.48 between percept duration and GABA to be detected with a power of 0.80. Participants were recruited from advertisements placed within the University of Melbourne and Monash University. Eligible participants in the migraine group were required to have been formally diagnosed by a general practitioner or neurologist and/or have symptoms consistent with the International Classification of Headache Disorders (ICHD)-III criteria for migraine with or without aura (37). As part of the screening process, participants with migraine provided details of their migraine including age of first migraine onset, attack frequency, number of days since their last migraine, location of pain on the head, type and level of pain, and sensory symptoms/ aura (Table 1). Individual migraine severity was graded based on the Migraine Disability Assessment (MIDAS). Eligible non-headache controls were required to have never experienced a migraine, have not suffered from any unexplained headaches and have not experienced more than five spontaneous headaches such as those arising from sickness and tension. All participants had uncorrected or corrected vision of ±5D spherical and ±2D cylindrical with a resultant visual acuity of no worse than 6/7.5. Participants did not suffer from any other visual or neurological conditions and were also not taking any regular medications known to affect vision and cognition, including migraine prophylactics. Prior to the commencement of testing, all participants provided written informed consent as approved by the Human Research Ethics Committee of Monash University (MUHREC CF13/2885-2013001549) and in accordance with the Declaration of Helsinki. All participants completed all parts of the study. Participant recruitment and data collection were conducted between December 2015 and July 2016.

**Table 1:**
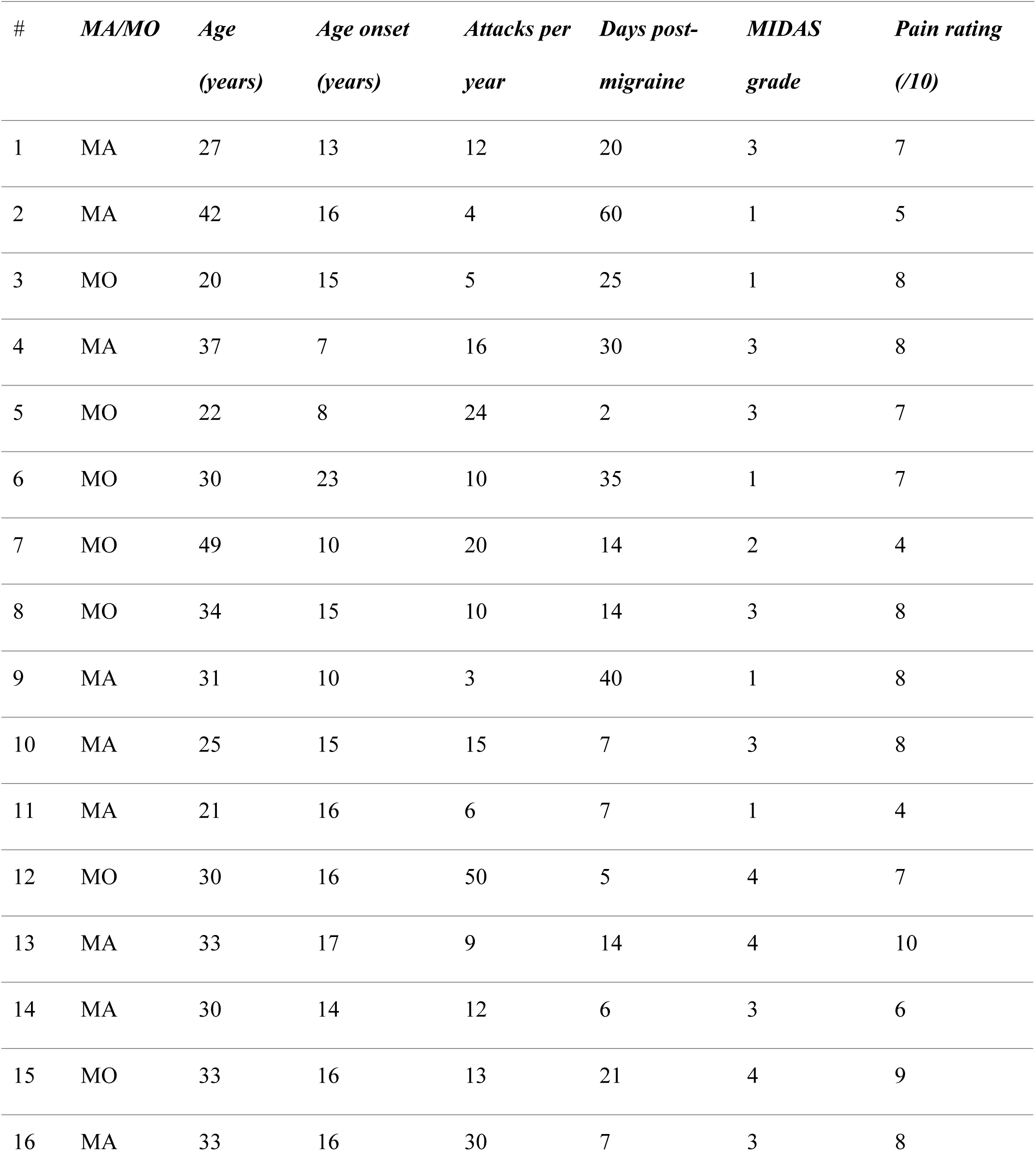
Demographics of the migraine group including whether individuals were aura or non-aura sufferers (MA/MO), age at time of testing, age of first migraine onset, number of attacks per year, number of days post-migraine at time of testing, MIDAS grade and average pain rating of their migraine attacks.

### 2.2 Magnetic resonance spectroscopy acquisition

All participants were positioned in a 3T MRI scanner (MAGNETOM Skyra, Siemens Healthineers, Erlangen, Germany) with a 32-channel head coil to collect the T_1_-weighted whole brain image (MPRAGE, repetition time [TR] = 58ms; echo time [TE] = 3.7ms, 1 mm^3^ isotropic voxels) and single voxel spectroscopy data (TR = 1500ms; TE = 68ms; voxel size = 30 × 25 × 20mm^3^). The visual cortex voxel was individually adjusted to be centred midline on either side of the calcarine sulcus and with 6mm anterior to the dura. This voxel placement was chosen to be consistent with previous work that has identified a relationship between occipital cortical GABA and binocular rivalry percept duration (33). Voxels were carefully positioned to avoid major blood vessels, meninges and ventricles. A prototype GABA-specific sequence of Point Resolved Spectroscopy (PRESS) with a previously described MEGA suppression scheme was used to acquire the 1H J-difference spectra (192 transients, each consists of two TRs; total scan time = 9 minutes) (36). A frequency selective inversion pulse of 1.9ppm was applied to the ^4^CH_2_ resonance of GABA during the odd transients (EDIT ON) while a pulse of 7.46 ppm was applied during the even transients (EDIT OFF). GABA concentration (institutional units (iu)) was quantified as the difference between the EDIT ON and EDIT OFF spectra. Given that the edited GABA signal is contaminated by co-edited macromolecules, GABA values are referred to as GABA+ in the rest of the paper. The unsuppressed water signal (8 averages) was acquired from the same voxel. GABA fit errors were used to assess the quality of spectra. All of the spectra had a GABA fit error < 10% (see Fig 1a for an example spectrum). Fit quality metrics (mean±SD: Cr fit error: 10.37±3.16; Cr FWHM: 7.85±0.40; GABA fit error: 6.75±0.68; GABA FWHM: 18.03±1.01; GABA signal-to-noise ratio: 14.98±1.48; Glx fit error: 6.12±0.97; Glx FWHM: 14.79±1.35; Glx signal-to-noise ratio: 16.72±2.52) were comparable to previous studies (31).

**Fig 1:**
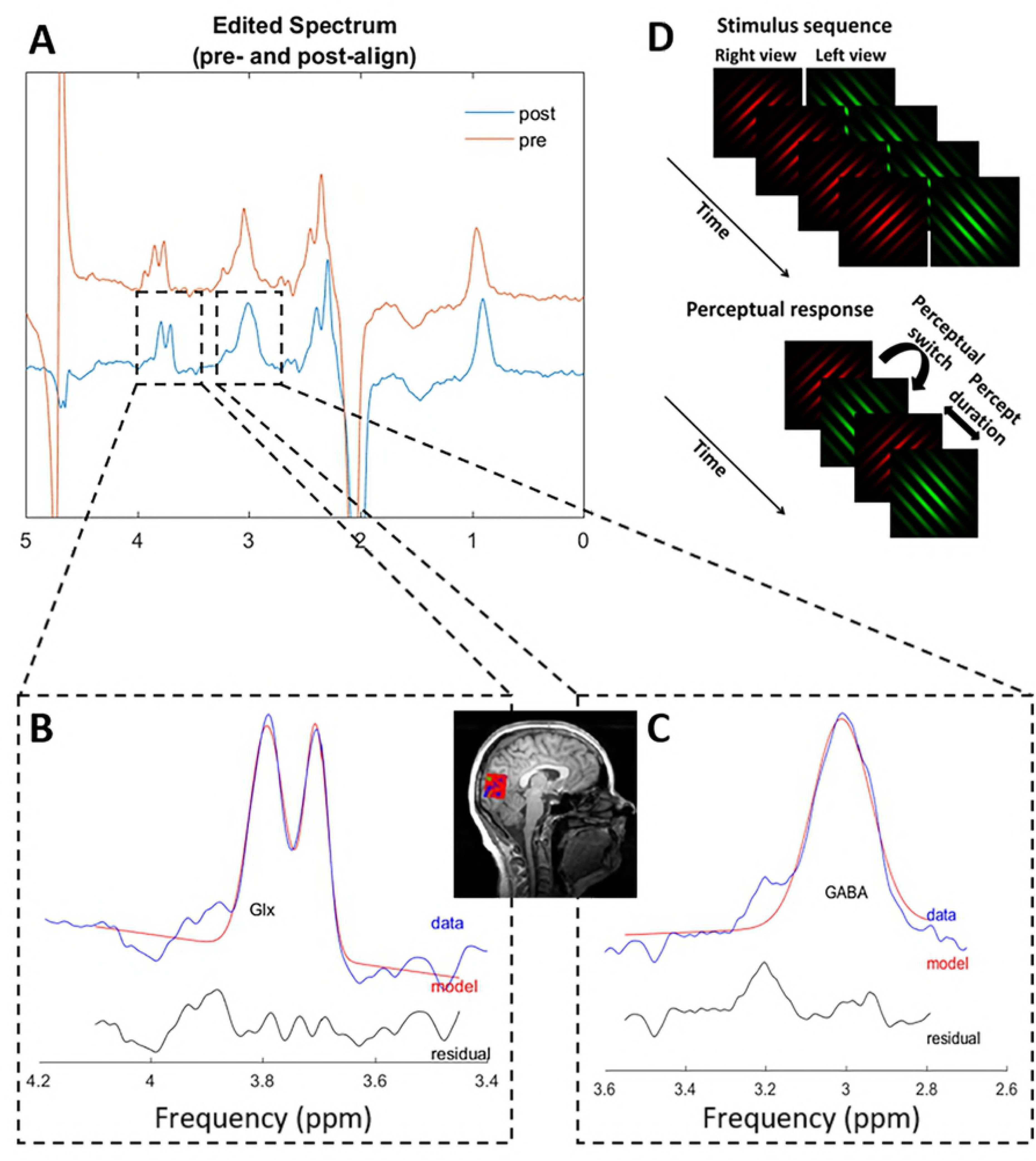
Example magnetic resonance spectroscopy spectrum and stimulus sequence for the binocular rivalry task. (a) An example spectrum measured from the visual cortex of a migraine participant. Frequency and phase correction are performed on the pre-aligned raw spectrum (red) using Gannet to generate a post-align spectrum (blue). Gannet then generates best fit curves of the Glx (b) GABA (c) peaks on the post-align spectrum (blue). Glx and GABA concentrations are defined as the area under the fitted curves (red). Inset figure illustrates the voxel placement at the occipital cortex. (d) Stimulus sequence used to assess binocular rivalry. A green grating oriented at 135° was shown to the left eye while a red grating oriented at 45° was shown to the right eye throughout each 90s test block. Observers perceived switches between the red and green grating within the 90s test blocks.

The GABA+ values were normalised to water (GABA+/water) and adjusted for tissue composition of the voxel using the Gannet software (38). Firstly, the T_1_-weighted images were segmented into grey matter (*GM*), white matter (*WM*) and cerebrospinal fluid (*CSF*) using the SPM8 software. GABA+ values were then corrected (*C*_*fullcorr*_) using the following equation proposed by Harris et al. (39):

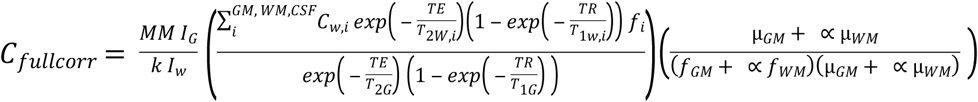

where, *MM* is the correction factor for co-edited macromolecular signal; I_G_ and I_W_ are the GABA and water signal integrals; c_w_ is the visible water concentration; *k* is the editing efficiency of GABA; *T*_1*G*_ *T*_1*w*_ *T*_2*G*_ *T*_2*w*_ are the T1 and T2 relaxation time constants for GABA and water; *TE*: echo time; *TR*: Repetition time. *f*_*GM*_ and *f*_*WM*_: grey matter and white matter volume fractions; μ_*GM*_ and μ_*WM*_: control group average grey matter and white matter fractions; ∝: assumed ratio between GABA concentrations in grey matter and white matter. The equation and the values for *MM*, *k*, **C**_*w*_, *T*_1*G*_, *T*_1*w*_, *T*_2*G*_,*T*_2*w*_, and ∝ were taken from GannetQuantify routine from Gannet toolbox (Fig 1c). The scan sequence was optimised for the study of GABA but we were also able to obtain an estimate of the excitatory glutamate neurotransmitter in the form of a glutamate-glutamine complex (edited-Glx). Edited-Glx was estimated using the Gannet software and normalised to water (Edited-Glx/water is expressed as institutional units (iu)) (Fig 1b). The normalised value was further corrected for voxel CSF-fraction volume differences using the following equation:

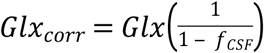

where, *Glx*_*corr*_ is CSF-fraction-corrected edited-Glx/water, *Glx* is the raw edited-Glx/water value in arbitrary units, and *f*_*CSF*_ is the voxel’s CSF fraction.

### 2.3 Binocular rivalry task

Participants performed a binocular rivalry task immediately after the MRI scan in a separate room. Stimuli were presented on a gamma-corrected Sony G520 21-inch CRT monitor (refresh rate 120Hz, 800 × 600 pixels, maximum luminance 100cd/m^2^) via a ViSaGe graphics system (Cambridge Research Systems, Kent, UK) using custom software written in Matlab (version 7, The MathWorks.inc., USA). The stimuli consisted of two circularly windowed sine gratings of 4° diameter and 1.5 c/deg spatial frequency (Fig 1d). One Gabor contained black-green (CIE x,y = 0.30, 0.59) gratings oriented at 135 degrees while the other patch contained black-red (CIE x,y = 0.58, 0.35) gratings oriented at 45 degrees, presented on a uniform grey background of 35cd/m^2^. Participants viewed the stimuli binocularly through a mirror stereoscope at a viewing distance of 53cm such that the right eye viewed a red grating oriented at 45° whereas the left eye viewed a green grating oriented at 135°. Perceptually, this stimulus will regularly switch from being perceived as a red grating to a green grating every few seconds. All participants were given a practice run and then performed four runs each of 90 seconds duration in which they indicated their percept by pressing the left button for a dominant green grating, right button for red grating and the middle button for a mixed red-green percept. Individual thresholds on this task were determined by the median percept duration of the red-only and green-only percepts.

### 2.4 Statistical analysis

Statistical analyses were conducted in SPSS Statistics Version 21 (IBM, New York, USA). Repeated measures Analysis of Variance (RM-ANOVA) and post-hoc independent sample t-tests were used to assess group differences in spectroscopy measured neurotransmitter levels (within group factor: neurotransmitter, between group factor: participant group). When a normality test failed, groups were compared using a Mann-Whitney Rank Sum Test. Pearson’s correlations were conducted to assess the relationships between the spectroscopy measures, perceptual measures and the number of days since individuals’ last migraine attack.

## 3 Results

### 3.1 Differences in neurochemical concentration between groups

There was no main effect of participant group on the spectroscopy measured neurotransmitters (RM-ANOVA: F(1,30)=2.81, *p* = 0.10) but there was a significant interaction between neurotransmitter type and group (F(1,30)=8.98, *p* = 0.005). Post-hoc t-test revealed that migraineurs had similar occipital GABA+/water level to controls (mean±sd: Controls: 3.31±0.32iu, Migraineurs: 3.42±0.24iu; t-test: t(30)=1.12, *p* = 0.27) (Fig 2a). However, the edited-Glx/water estimate was significantly reduced in migraineurs relative to controls (Controls: 1.90±0.27iu, Migraineurs: 1.61±0.12iu; Mann-Whitney Rank Sum Test: U = 36.00, *p* < 0.001) (Fig 2b), resulting in a significantly higher GABA/Glx ratio in the migraine cohort (Controls: 1.78±0.36iu, Migraineurs: 2.14±0.19iu; t(30)=3.54, *p* = 0.001) (Fig 2c).

**Fig 2:**
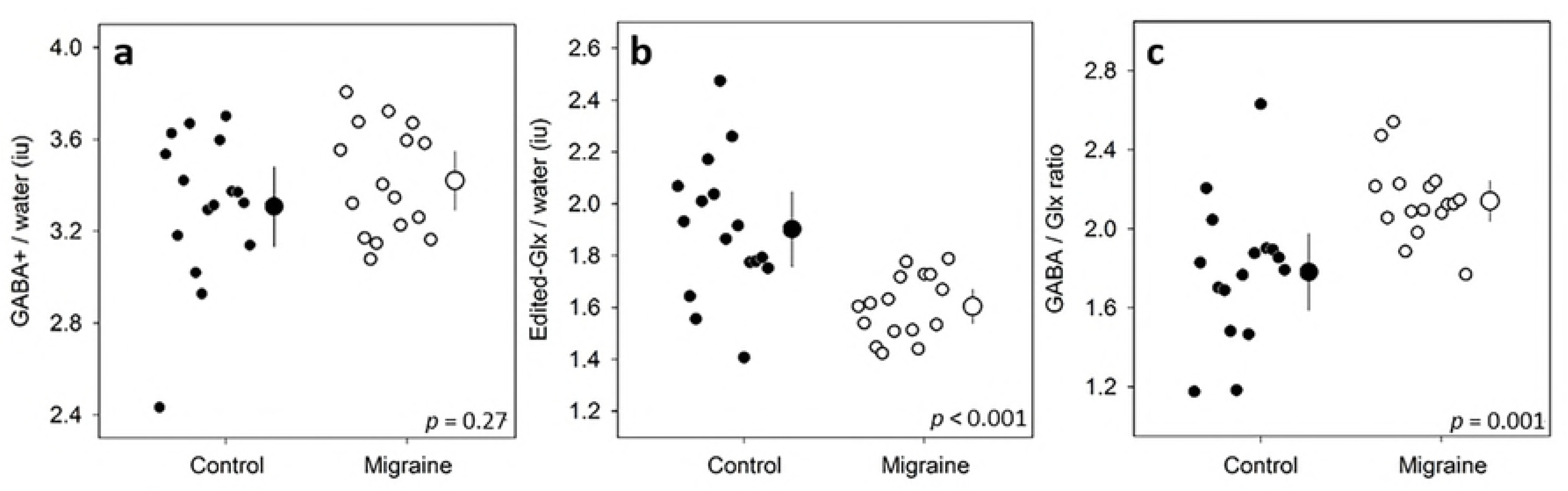
Group averaged GABA and Glx levels. Individual measures (small symbols, jittered along the x-axis) and group averages (large symbols; error bars represent 95% confidence intervals) of occipital GABA+/water (a), Edited-Glx/water (b) and GABA/Glx ratio (c) obtained from the visual cortex of controls (filled) and migraineurs (unfilled).

### 3.2 No difference in perceptual rivalry rate between groups

Perceptual performance for the binocular rivalry task was compared based on median percept duration of the red-only and green-only percepts (Fig 3). On average, those with migraine had a median perception duration of 2.08s (±0.66s), which was not statistically different (U=101.00, *p* = 0.47) from that measured in the control group (1.85±0.73s). Rivalry mixed percept duration was similarly not statistically different between migraineurs (median,IQR: 2.39,11.61s) and controls (0.00,5.19s) (Mann-Whitney U-test, U=88.00, *p* = 0.18) with a subset in each group reporting no mixed percepts at all (5 migraineurs, 10 controls).

**Fig 3:**
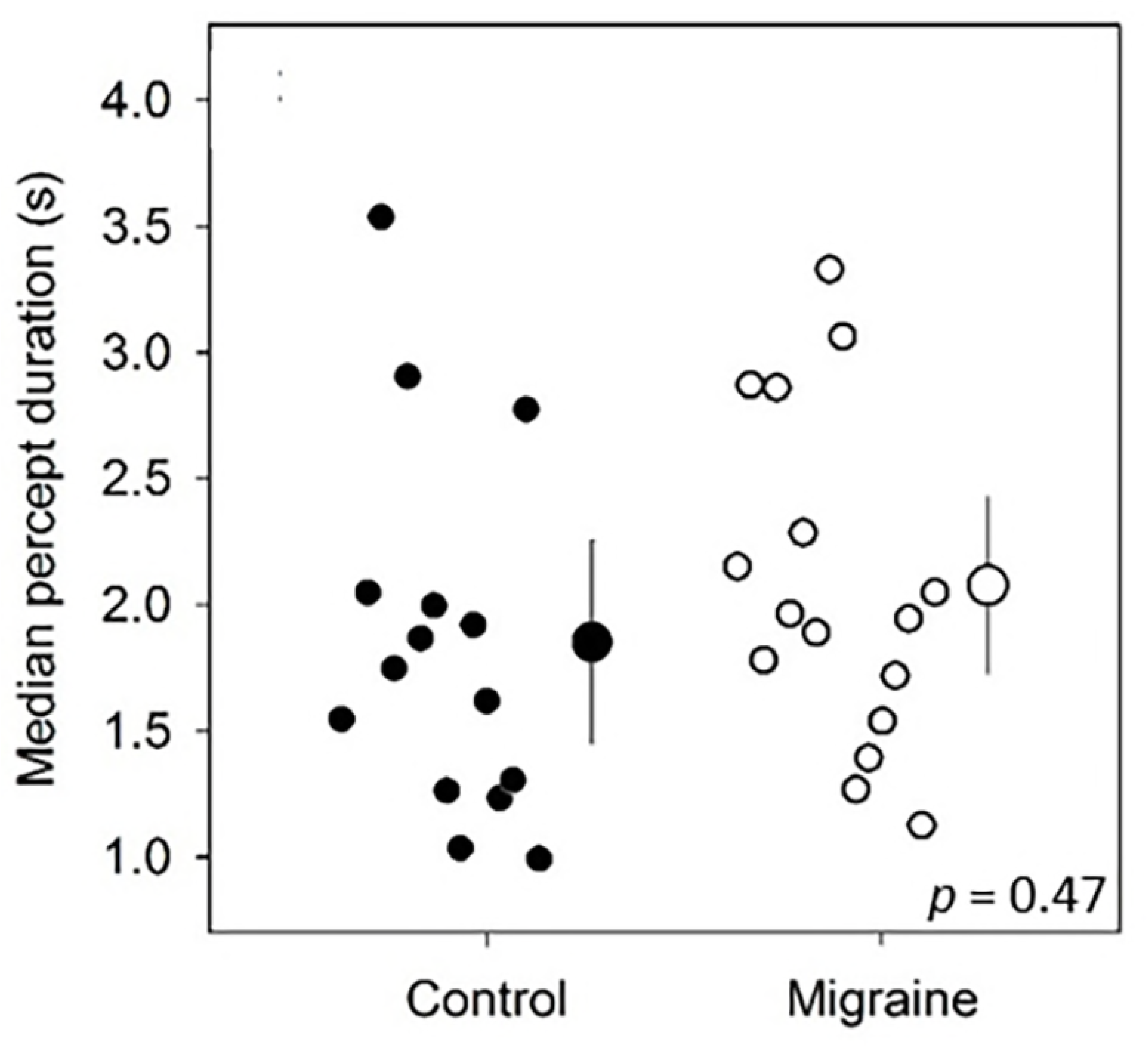
Group averaged rivalry percept duration. Individual measures (small symbols, jittered along the x-axis) and group averages (large symbols; error bars represent 95% confidence intervals) of median percept duration (a) in controls (filled) and migraineurs (unfilled).

### 3.3 No relationship between perceptual rivalry rate and neurochemical concentration

A significant positive correlation between GABA and percept duration estimates has been previously reported in healthy younger adults (33), therefore we assessed if the same trend holds in this dataset. A Pearson’s correlation analysis performed on the entire dataset (both the controls and migraineurs), did not reveal a significant relationship between these measures (r = 0.23, *p* = 0.21) (Fig 4).

**Fig 4:**
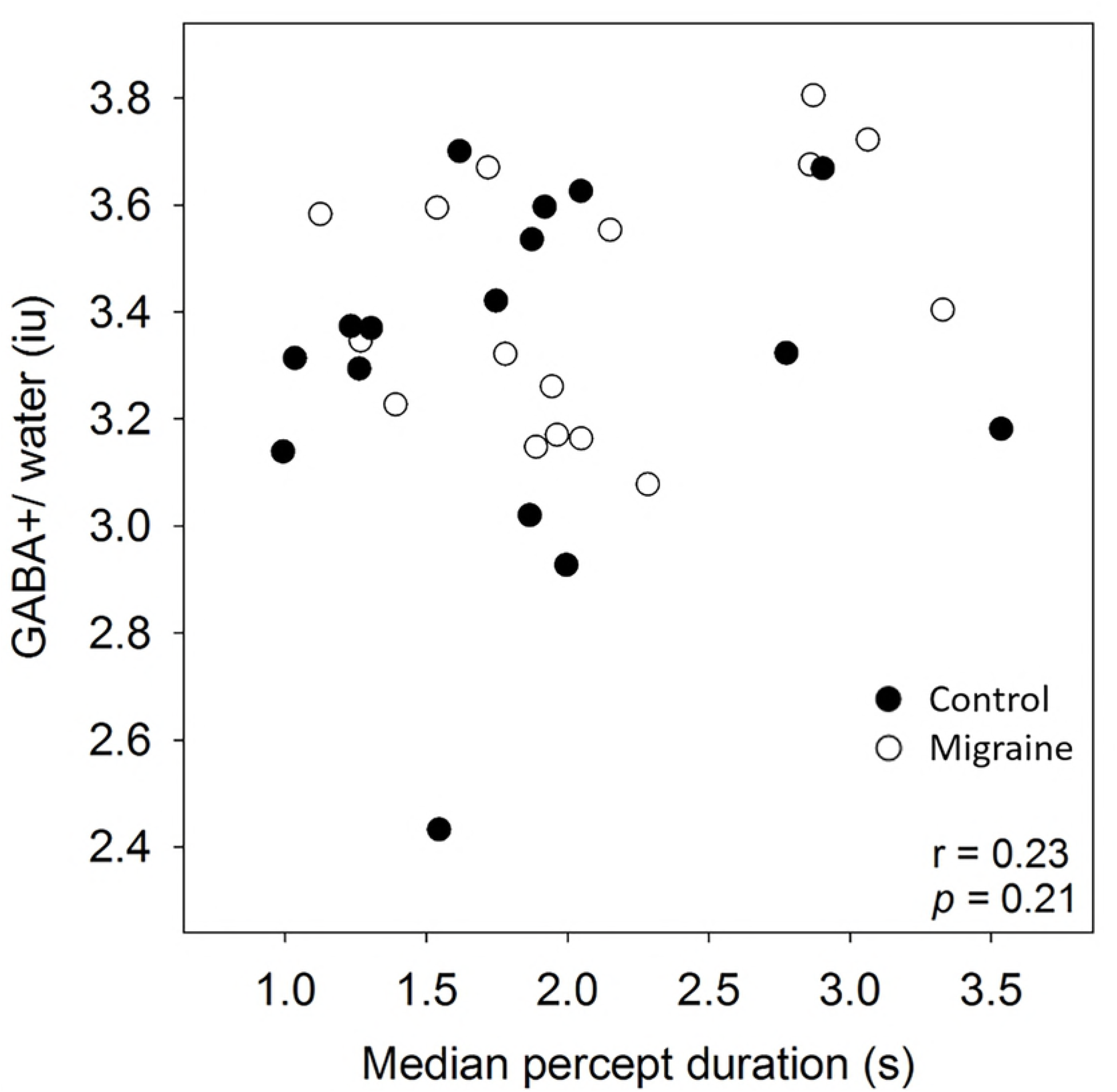
Relationship between median percept duration for the binocular rivalry task and the GABA+/water in visual cortex. Data is shown for each group separately but pooled to determine the correlation.

## Discussion

The aims of this study were to determine if: (1) occipital cortex inhibitory (GABA+) or excitatory (Glx) levels differ between those with migraine and controls during the non-headache period, and (2) whether levels of these neurochemicals can explain any of the variability in perceptual binocular rivalry percept duration. Our data reveals differences in the estimate of edited-Glx/water between groups, and in the related balance between GABA+/Glx ratio, but not in GABA+ *per se* (Fig 4). Contrary to previous reports (26, 27), there was no group difference in binocular rivalry percept duration (Fig 4), nor was there a correlation between percept duration and occipital cortex GABA+ (Fig 4).

Two previous studies have reported slower perceptual switch rates (i.e. longer percept duration) in those with migraine than controls (26, 27): a finding that we did not replicate here (Fig 4). Wilkinson et al. (2008) used a similar binocular rivalry stimulus but that was monochromatic, subtended a smaller visual angle and contained gratings of a higher spatial frequency (26), all of which could potentially influence the perceptual estimates. The other study did not use binocular rivalry, but instead measured perceptual rivalry using a drifting plaid stimulus, in addition to an auditory rivalry stimulus (27). The neural processing of both stimulus types involves brain regions outside of the primary visual cortical voxel measured here (27). While previous work indicates a role for GABA-ergic inhibition in mediating binocular rivalry strength (31, 33), it is likely that a range of other neuromodulators can also impact rivalry dynamics, such as serotonin (35, 40) and noradrenaline (41).

Our migraine group had similar average levels of GABA in the occipital region as compared to non-headache controls (Fig 4). We used a scan sequence (MEGA-PRESS) more specific for quantifying GABA than previous studies, however, similar to previous work, did not identify any difference in mean GABA levels between groups (11). Our study sample size was chosen to have a power of 0.8 to find a correlation between binocular rivalry and GABA+ of similar size to that reported by van Loon et al. (2013) (r = 0.51). A recent study has confirmed the relationship between binocular rivalry percept duration and occipital cortex GABA+ in healthy adults, however the relationship was weaker (r = 0.35) (31). Our data is consistent with the idea that any relationship between perceptual rivalry duration and GABA+ in visual cortex, if present, is weak.

In contrast to previous reports of elevated excitatory Glx in people with migraine (12, 13), and inconsistent with Bridge’s finding for no difference in occipital Glx levels (10), we found reduced occipital Glx levels in the migraine group as compared to the non-headache controls (Fig 4). Participant demographics were markedly different between studies such as limited to people who experience migraine with aura (13) and limited to only female observers (10, 12), whereas we included participants experiencing either migraine with and without aura of both genders. As migraine is a cyclic condition, the time pre-post migraine at which patients are being tested for Glx and GABA might influence the measured levels. It is unclear from two of these previous studies how many days post-migraine each observer was when tested with reports only describing that participants were at least 1 day post-migraine (12) or 5 days (13) pre-post migraine. However, it is important to note that the scan sequence applied in our experiment was optimised to quantify GABA, so caution should be taken when interpreting measurement outcomes of other cortical metabolites quantitatively, including Glx.

Our experiments were motivated by previous reports of data collected from healthy younger (33) and older (31) controls that suggested a probable relationship between occipital GABA levels and rivalry percept duration. However, in our sample, we did not find a correlation between percept duration and any of the neurochemical concentrations measured. Of course, occipital inhibitory function is not the only mechanism underlying binocular rivalry (42).

Alterations to serotonin modulation have been proposed previously to impact on the rate of rivalry alternations (26, 34, 40). Evidence from human pharmacological studies report observers having slower switch rates (longer percept duration) after being administered with serotonin-agonists that when bound to its receptor, shuts down further release of serotonin (34, 40). This is consistent with a proposed mechanism for downregulated serotonergic function interictally (16, 17). Low serotonin levels are known to predispose several migraine triggers like stress (15) and to contribute to migraine pain via the trigeminovascular nociceptive pathway (43). Integrating these two findings, it is possible that we would measure lower serotoninergic function with slower rivalry switch rates in individuals just before a migraine event. However, in the absence of any formal measurement of occipital serotonin levels and without longitudinal data followed in the same individuals, we can only speculate regarding the relationship between serotonin and rivalry percept in migraine.

In summary, our data revealed no difference in GABA levels but reduced excitatory cortical neurotransmitter concentration in the occipital cortex of people with migraine in between their attacks, when compared to individuals without regular headaches. Despite previous reports of a correlation between GABA level and perceptual rivalry switching rates, in our study GABA levels and rivalry percept duration were not correlated, suggesting that GABA levels are not the primary driver of variance in perceptual rivalry in this situation. Our data also demonstrates that the previously reported relationship between rivalry percept duration and GABAergic inhibitory neurotransmitter concentration in visual cortex is not strong enough to enable rivalry percept duration to be assumed to be a surrogate for GABA concentration. Future work may also consider other plausible neuromodulators such as serotonin. Given the possible importance of duration pre-post migraine on perceptual measures and neurochemical status, significant insight might be gained from testing people with migraine at multiple timepoints within their migraine cycle.

## 5 Acknowledgements

This work was supported by the National Health and Medical Research Council (NHMRC) Project Grant (APP1081874).

## 6 Data availability

The datasets generated during and/or analysed during the current study are available as supplementary material.

